# Nanobodies against *Plasmodium* adhesins that block receptor engagement and malaria parasite invasion

**DOI:** 10.64898/2026.05.20.726696

**Authors:** Jaison D Sa, Jill Chmielewski, Amy Adair, Li Lynn Tan, Li-Jin Chan, Lucas Krauss, Kathleen Zeglinski, Quentin Gouil, John Chen, Colin Jackson, Christoph Q Schmidt, Sarel J Fleishman, Phillip Pymm, Wai-Hong Tham

## Abstract

Malaria is caused by *Plasmodium* parasites, and its clinical symptoms are a result of parasite invasion of red blood cells and the subsequent cycles of replication and proliferation. In human populations, *Plasmodium vivax* is responsible for the most widely distributed recurring malaria infections whereas *Plasmodium falciparum* inflicts the most mortality and morbidity. One well-characterized family of adhesins involved in red blood cell invasion is the reticulocyte-binding-like protein homolog family, known as the RBL superfamily which includes the PfRh family in *P. falciparum* and PvRBP family in *P. vivax*. Here we report a collection of nanobodies against three members of this adhesin family, PfRh5, PfRh4 and PvRBP2b. Nanobodies against these *Plasmodium* adhesins bind with high affinity across several epitopes and can block receptor engagement and inhibit parasite invasion of red blood cells. Using computational design, we generated stabilized PfRh4 variants that encompass the conserved scaffold present in the PfRh and PvRBP families of adhesins and show that several variants with improved expression retained binding to mouse monoclonal antibodies, nanobodies and Complement Receptor 1, the human receptor for PfRh4. We also observed that most of the inhibitory nanobodies against the three antigens recognized the conserved structural scaffold that define this family of adhesins. These results demonstrate the potential of nanobodies to block malaria parasite invasion into red blood cells.

## Introduction

Malaria parasites are exquisitely adapted for invasion and survival within human red blood cells. Parasite invasion begins with initial attachment of merozoites, the invasive form of malaria parasites, to red blood cells resulting in deformation of the red blood cell membrane [1,2]. The merozoite then orientates itself so that parasite adhesins at the apical tip are in direct contact with the red blood cell surface, allowing interaction with their cognate receptors to mediate irreversible attachment and commitment to invasion [3,4]. A tight junction is formed between the parasite and the red blood cell membrane [5] and active invasion proceeds as the tight junction moves from the apical to posterior pole of the merozoite. Once the merozoite is inside, the red blood cell membrane seals behind it completing invasion. The entire invasion process is completed within a few minutes [1]. The subsequent cycles of parasite growth, replication of parasites and egress from infected red blood cells are responsible for the symptoms associated with malaria [6].

Invasion of red blood cells by malaria parasites is governed by specific interactions between parasite adhesins and red blood cell receptors[7]. *P. falciparum* and *P. vivax* have two homologous families of parasite adhesins. The first is the erythrocyte-binding ligand family (EBL) in *P. falciparum* and its homologous family, Duffy-binding protein (DBP) family in *P. vivax* [8]. The second is the reticulocyte-binding homolog family in *P. falciparum* (PfRh) and its homologous family, the reticulocyte-binding protein family in *P. vivax* (PvRBP) [8]. PfRh and PvRBP are part of the Reticulocyte Binding Like (RBL) superfamily found in other *Plasmodium* species. In *P. falciparum*, the PfRh family consists of five members, namely PfRh1, PfRh2a, PfRh2b, PfRh4 and PfRh5[7]. In *P. vivax*, the PvRBP family consists of eight members, called PvRBP1a, PvRBP1b, PvRBP2a, PvRBP2b, PvRBP2c, PvRBP2d, PvRBP2e and PvRBP3 [9–11]. Several human receptors have been identified to bind to the PfRh and PvRBP family of proteins; PfRh5 to Basigin[12], PfRh4 to Complement Receptor 1 [13], PvRBP2b to Transferrin Receptor 1 [14,15] and PvRBP2a to CD98 [16].

There is a conserved homology region within the N-terminus of PfRh and PvRBP family of adhesins. While most PfRh and PvRBP proteins are large (230–350 kDa), PfRh5 represents one of the smallest members at 63 kDa. The crystal structure of PfRh5 provided the first structural insight to this conserved N-terminal domain, which shows a compact, primarily α-helical fold that is critical for binding its receptor basigin[17,18]. Similarly, structural studies of the N-terminal domains of PvRBP2a and PvRBP2b show high structural similarity to PfRh5, highlighting a conserved structural scaffold within the PfRh and PvRBP family of adhesins [14,15,19]. At present, there are no additional crystal structures of the other members of the PfRh and PvRBP family of proteins.

PfRh5 is a highly conserved and essential ligand for *P. falciparum* red blood cell invasion, making it a leading target for blood stage malaria vaccines [20,21]. Its interaction with basigin is critical for successful parasite invasion of red blood cells [12,17]. Recent studies characterized human monoclonal antibodies (mAbs) elicited by both vaccination and natural infection, providing insights into the antigenic landscape of PfRh5 and informing vaccine design strategies [22,23]. One study isolated and analyzed 186 human mAbs against PfRh5 from individuals either naturally exposed to malaria or vaccinated against it [23]. Most of these mAbs targeted non-neutralizing epitopes of PfRh5. However, two mAbs, MAD8-151 and MAD8-502 showed strong growth inhibitory activity by binding to PfRh5 near its basigin binding epitope[23]. These mAbs were structurally and genetically similar to highly effective vaccine-induced antibodies, sharing identical germline origins and binding orientations. Structural characterization confirmed that they block PfRh5-basigin interaction through steric hindrance. Of these, MAD8-151 significantly reduced parasitemia in humanized mice in *in vivo* experiments [23]. In another study, 236 human mAbs were isolated from individuals vaccinated with the RH5.1/AS01B vaccine [22]. Epitope mapping using surface plasmon resonance identified 12 distinct epitope communities. Representative mAbs from five of these communities were associated with parasite growth inhibitory activity by either directly blocking the PfRh5-basigin interaction or through steric hindrance [22]. Among these mAbs, R5.034 was the most potent inhibitor encoded by a recurrent public clonotype. It demonstrated the strongest parasite growth inhibitory activity, high affinity binding and its epitope did not overlap with any of the common PfRh5 polymorphisms [22].

PfRh4 is a red blood cell invasion ligand of *P. falciparum* and its interaction with Complement Receptor 1 (CR1) provides an alternate pathway for invasion of red blood cells that is distinct from sialic acid dependent pathways mediated by glycophorin receptors [13,24,25]. Several mouse mAbs against PfRh4 have been characterized but 5H12 is the only antibody identified to date with inhibitory activity [26]. 5H12 blocks complex formation between PfRh4 and CR1 as shown by fluorescence resonance energy transfer (FRET) and immunoprecipitation-based assays, suggesting that its epitope overlaps with the CR1 binding site on PfRh4 [26]. In addition, 5H12 inhibits *P. falciparum* parasite replication *in vitro*. At present, there are no structures for PfRh4 alone or in complex with antibodies, limiting structural insights into the inhibitory mechanism of 5H12 and the broader antigenic landscape of PfRh4.

PvRBP2b is a critical invasion ligand required for *P. vivax* entry into reticulocytes and it binds human Transferrin Receptor 1 (TfR1) in the presence of its endogenous ligand transferrin (Tf) [14,15]. Four inhibitory mouse mAbs 3E9, 4F7, 6H1 and 10B12 have been identified against PvRBP2b [14]. All four inhibit the interaction of recombinant PvRBP2b to TfR1 on the reticulocyte surface, as determined using a reticulocyte binding assay [14]. 3E9, 6H1 and 10B12 also block *P. vivax* invasion of reticulocytes in short-term *P. vivax ex vivo* invasion assays using Brazilian and Thai clinical isolates [14]. Structural studies revealed that these antibodies recognize distinct epitopes at the N-terminal domain between residues 214 and 416 [15]. 3E9 binds to the side of the N-terminal domain, interfering sterically with both TfR1 and transferrin (Tf). In contrast, 4F7, 6H1 and 10B12 bind near the tip of the N-terminal domain of PvRBP2b, with 4F7 and 6H1 sharing an overlapping epitope [15]. In addition to these mouse mAbs, several anti-PvRBP2b human mAbs derived from naturally exposed individuals have been characterized[27]. Eight of these mAbs namely 239229, 241242, 253245, 258259, 260261, 262231, 273264 and 346343 strongly inhibited PvRBP2b-TfR1-Tf complex formation[27]. These eight mAbs also inhibited PvRBP2b binding to reticulocytes with > 86 % inhibition. Epitope mapping revealed that the majority of these mAbs recognized epitopes within the N-terminal domain of PvRBP2b between residues 166 and 450 [27]. Two mAbs, 335338 and 340341 were classified as intermediate inhibitors. Interestingly, one mAb 297280 enhanced PvRBP2b binding to reticulocytes.

Together, these studies highlight the critical roles of PfRh5, PfRh4 and PvRBP2b in red blood cell invasion and showcase how monoclonal antibodies can effectively block receptor engagement and inhibit parasite invasion. For this study, we characterize the first collection of nanobodies against these three *Plasmodium* adhesins, PfRh5, PfRh4 and PvRBP2b, and show that they can block receptor engagement and inhibit parasite invasion. In addition, we show that most inhibitory nanobodies bind to the conserved structural scaffold that is present in the homologous PfRh and PvRBP family of adhesins.

## Results

### Isolation of PfRh5, PfRh4 and PvRBP2b specific nanobodies

To isolate nanobodies targeting PfRh5, we immunised two alpacas with recombinant PfRh5 and generated the associated phage display libraries. Two rounds of phage display panning were performed and nanobody sequences were identified by next-generation sequencing (NGS) and Sanger sequencing. NGS sequences of nanobodies were clustered based on 90% similarity in their complementary determining region 3 (CDR3) using Alpseq[28] (Figure 1A). From the NGS data, we selected the top 16 most abundant nanobodies after two rounds of phage display selection (2886 – 2901) (Data S1). We identified eight nanobodies (2902 – 2909) from Sanger sequencing with an enrichment score of two or more (Figure S1, Data S1). Using biolayer interferometry (BLI), all 24 PfRh5 nanobodies bound to recombinant PfRh5 with nanomolar affinity with *K*D values ranging from 1.9 to 10.4 nM (Figure 1B, Data S1). Epitope binning experiments showed that PfRh5 nanobodies separated into five major epitope bins (Group A – E, Figure 1C, Figure S1). Group A represents the largest group with 11 nanobodies and the smallest group is Group C with one nanobody 2906 (Figure 1C). Group B includes two nanobodies, 2890 and 2900. The eight nanobodies in Group D are 2887, 2889, 2894, 2899, 2901, 2902, 2903 and 2907, and Group E has two nanobodies, 2895 and 2904.

**Figure 1.**
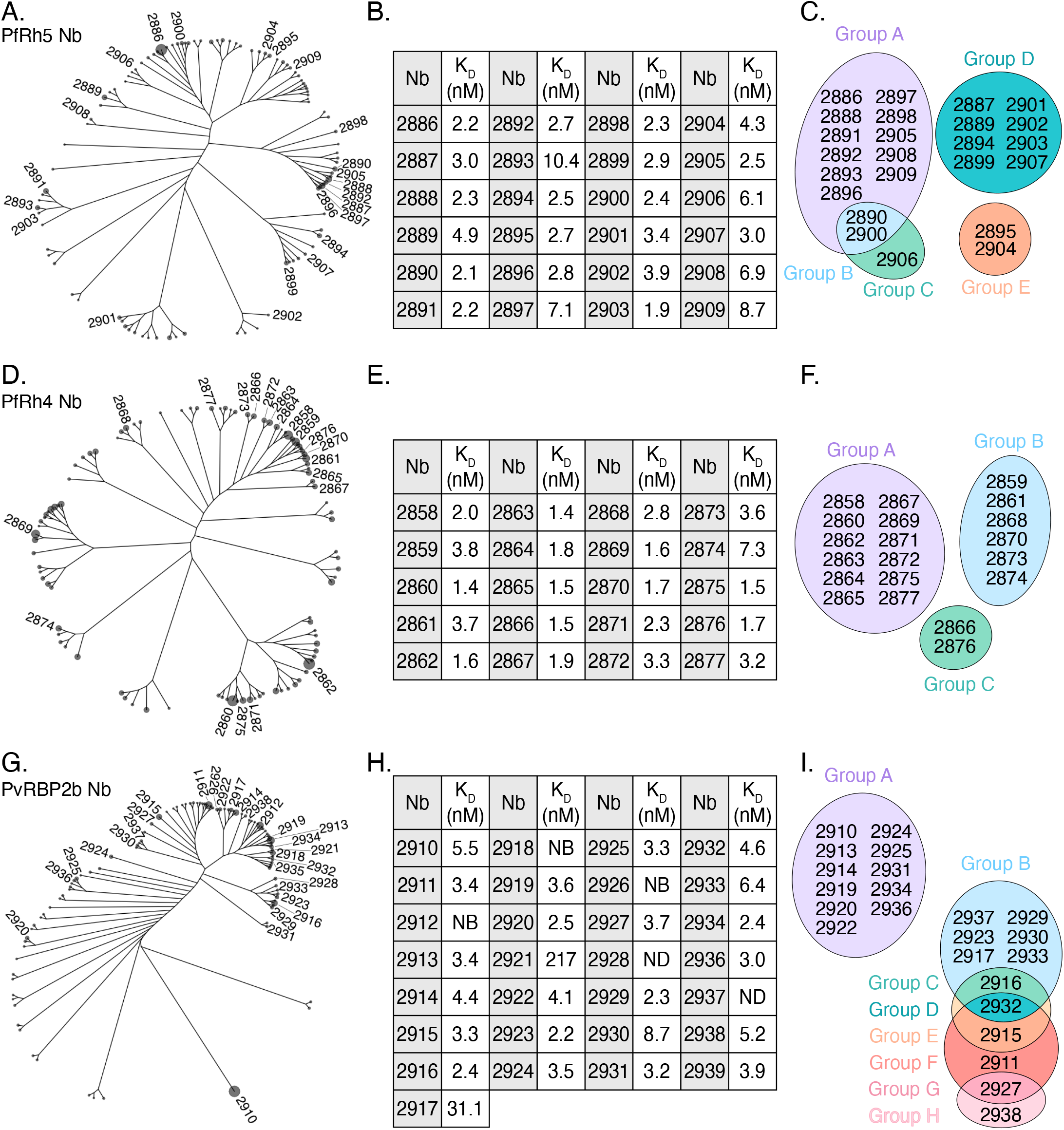
Identification and characterization of PfRh5, PfRh4 and PvRBP2b specific nanobodies. Cladograms showing the 100 most abundant nanobodies (Nb) from next-generation sequencing (NGS) of round 2 phage display libraries panned against PfRh5 (A), PfRh4 (D) and PvRBP2b (G). Nanobodies selected for further characterization are labelled, with branch tips scaled relative to abundance (counts per million). Binding affinities (*K*_D_) of selected nanobodies for PfRh5 (B), PfRh4 (E) and PvRBP2b (H), determined by bio-layer interferometry (BLI). NB represents no binding, and ND represents not determined due to low affinity. Values represent means from two independent experiments (n=2). Venn diagrams summarizing epitope binning results for PfRh5 (C), PfRh4 (F) and PvRBP2b (I) nanobodies from competition experiments using BLI. Epitope groups A–H are color-coded. Nanobodies in the same partition represent competition for the same epitopes.

To generate nanobodies against PfRh4, we immunised an alpaca with recombinant PfRh4 and generated the associated phage display library. For nanobody discovery analyses, we used similar approaches as for the PfRh5 nanobodies above. From the NGS data, we selected the top 20 most abundant nanobodies after two rounds of phage display (2858 - 2877) which were also present in the Sanger sequencing results (Figure 1D, Data S1). All 20 PfRh4 nanobodies bound to recombinant PfRh4 with nanomolar affinity with *K*_D_ values ranging from 1.4 to 7.3 nM (Figure 1E, Data S1). PfRh4 nanobodies separated into three major epitope bins (Group A - C, Figure 1F, Figure S1). Group A represents the largest group with 12 nanobodies. The six nanobodies in Group B are 2859, 2861, 2868, 2870, 2873 and 2874. The smallest group is Group C with two nanobodies 2866 and 2876.

For PvRBP2b, from the NGS data, we selected the top 29 most abundant nanobodies after two rounds of phage display (2910 - 2938) (Figure 1G, Data S1). Among these, 24 nanobodies were also present in Sanger sequencing results (Data S1). We identified one nanobody (2939) from Sanger sequencing which was absent in the top 100 most abundant nanobodies from the NGS dataset (Data S1). Using BLI, we observed that 2912, 2918, and 2926 did not bind recombinant PvRBP2b, and 2928 and 2937 affinities were not determined due to low affinity. The other 24 nanobodies bound with *K*_D_ values ranging from 2.2 to 217.0 nM (Figure 1H, Data S1). PvRBP2b nanobodies separated into eight epitope bins (Group A - H, Figure 1I, Figure S1). Group A represents the largest group with 11 nanobodies, followed by Group B with six nanobodies, 2937, 2923, 2917, 2929, 2930, and 2933. Groups C through H had one nanobody each, with Group C and D nanobodies, 2916 and 2932, overlapping with Group B nanobodies, and Groups C through H nanobodies showing some binding overlap. For this epitope binning assay, we used the N-terminal structural scaffold domain, PvRBP2b_169-470_ and since nanobodies 2921, 2928 and 2939 bind outside of this domain, they were not included in the epitope binning experiment (Data S2A).

Collectively these results show that phage display campaigns identified high affinity nanobodies against their target *Plasmodium* adhesins. These nanobodies bound in three to eight different epitope groups. We also visualised the sequence space of the selected nanobodies against the three *Plasmodium* adhesins and observed that nanobodies enriched toward different *Plasmodium* adhesins can occupy overlapping parts of sequence space, but hotspots of highest enrichment are distinct when targeting different adhesins (Figure S2).

### Nanobodies against *Plasmodium* adhesins block complex formation with their human receptors

To determine whether these nanobodies against the *Plasmodium* adhesins could block receptor engagement, we established FRET-based assays for the three pairs of adhesin-human receptor interactions; basigin-PfRh5, CR1-PfRh4 and TfR1-Tf-PvRBP2b. FRET signal in the absence of nanobody was designated as 100 % FRET signal (no Nb, Figure 2A - C). The 0 % FRET intensity was arbitrarily defined based on addition of 1 % SDS, which denatures the proteins and prevents complex formation (SDS, Figure 2A - C). Based on % FRET intensity, we categorized strong inhibitors (< 20 % FRET intensity, green), intermediate inhibitors (20–80 % FRET intensity, yellow), non-inhibitors (> 80 % FRET intensity, blue) and enhancers (> 150% FRET intensity, purple). As a negative control, we used a SARS-CoV-2 nanobody, WNb27, which did not decrease the FRET intensity (Figure 2A - C)[29].

**Figure 2.**
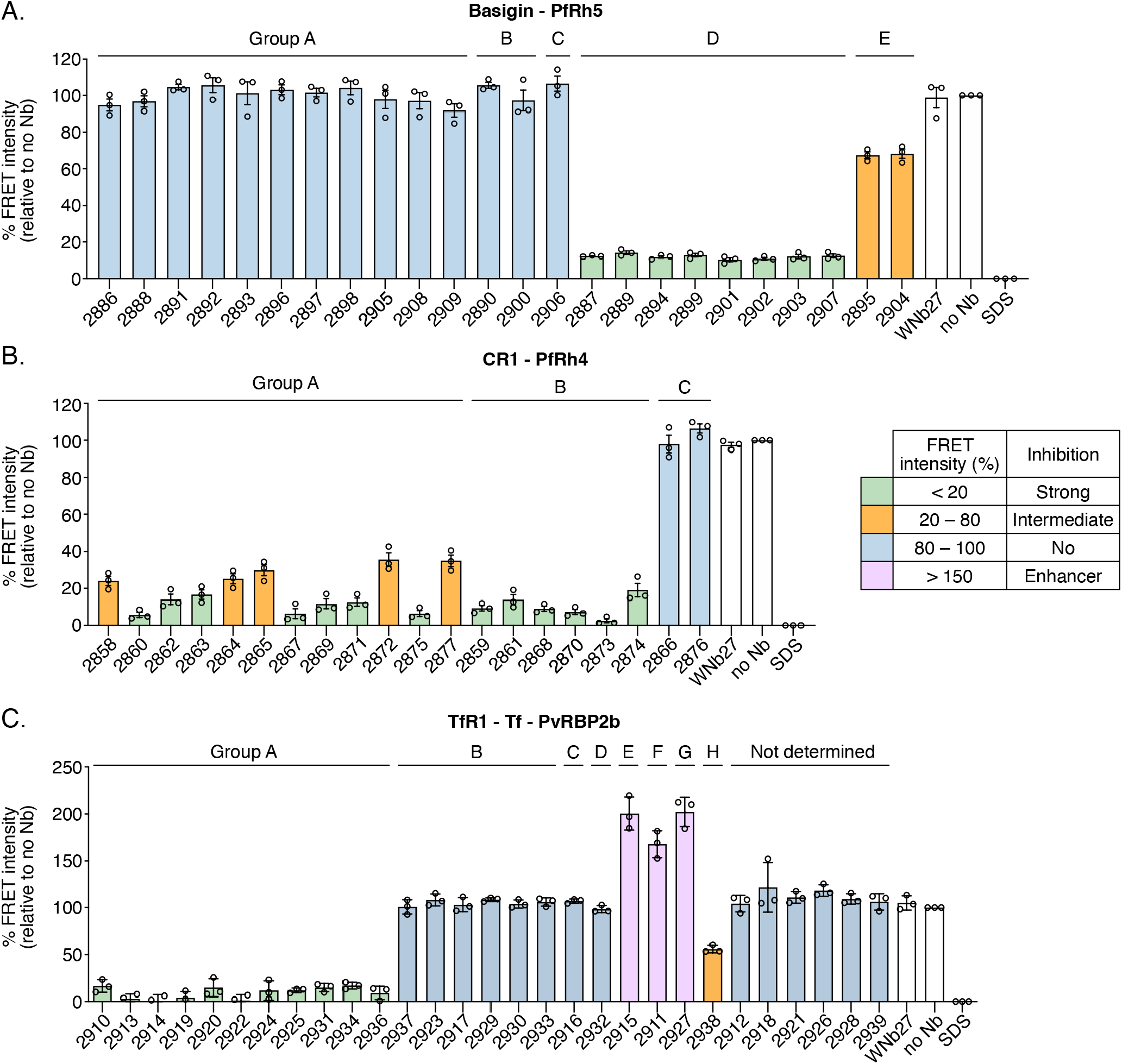
PfRh5, PfRh4 and PvRBP2b nanobodies that block complex formation. FRET-based assays were used to analyze nanobody inhibition of (A) PfRh5-basigin interaction, (B) PfRh4-CR1 interaction and (C) PvRBP2b-TfR1-Tf interaction. FRET intensity in the absence of nanobodies was defined as 100%. Data are presented as mean ± SEM from three independent experiments (n=3), with each circle representing one independent experiment.

Using this assay, we observed that all eight PfRh5 nanobodies from Group D were strong inhibitors (green, Figure 2A). Both PfRh5 nanobodies from Group E, 2895 and 2904, were intermediate inhibitors (yellow, Figure 2A). In contrast, nanobodies from Group A, Group B and Group C were non-inhibitors (blue, Figure 2A). For PfRh4 nanobodies, seven from Group A, 2860, 2862, 2863, 2867, 2869, 2871 and 2875 and all six PfRh4 nanobodies from Group B were strong inhibitors (Figure 2B). The remaining five PfRh4 nanobodies from Group A, 2858, 2864, 2865, 2872 and 2877 were intermediate inhibitors (yellow, Figure 2B). In contrast, the two PfRh4 nanobodies from Group C were non-inhibitors (blue, Figure 2B). For PvRBP2b nanobodies, all Group A nanobodies were strong inhibitors and the single nanobody 2938 in Group H was an intermediate inhibitor (Figure 2C). It is noted that all Group A nanobodies against PvRBP2b also recognize the conserved structural scaffold present in the N-terminal domain (PvRBP2b_169-470_, Data S2). While the other nanobodies did not inhibit complex formation, nanobody 2915 from Group E, nanobody 2911 from Group F and nanobody 2927 from Group G appear to increase the FRET signal which suggests that they enhance complex formation (Figure 2C).

### Nanobodies against PfRh5 and PfRh4 inhibit *Plasmodium falciparum* parasite invasion

We wanted to determine if the PfRh5 and PfRh4 nanobodies could block invasion using the parasite growth inhibition assays. Parasite growth was measured using flow cytometry following one replication cycle, and 100 % growth was defined as the parasitemia (percentage of all red blood cells infected) in the absence of nanobody (no Nb, Figure 3A, B). In the following growth assays, we describe these categories: strong inhibitors < 20 % growth, green), intermediate inhibitors (20–80 % growth, yellow) and non-inhibitors (> 80 % growth, blue).

**Figure 3.**
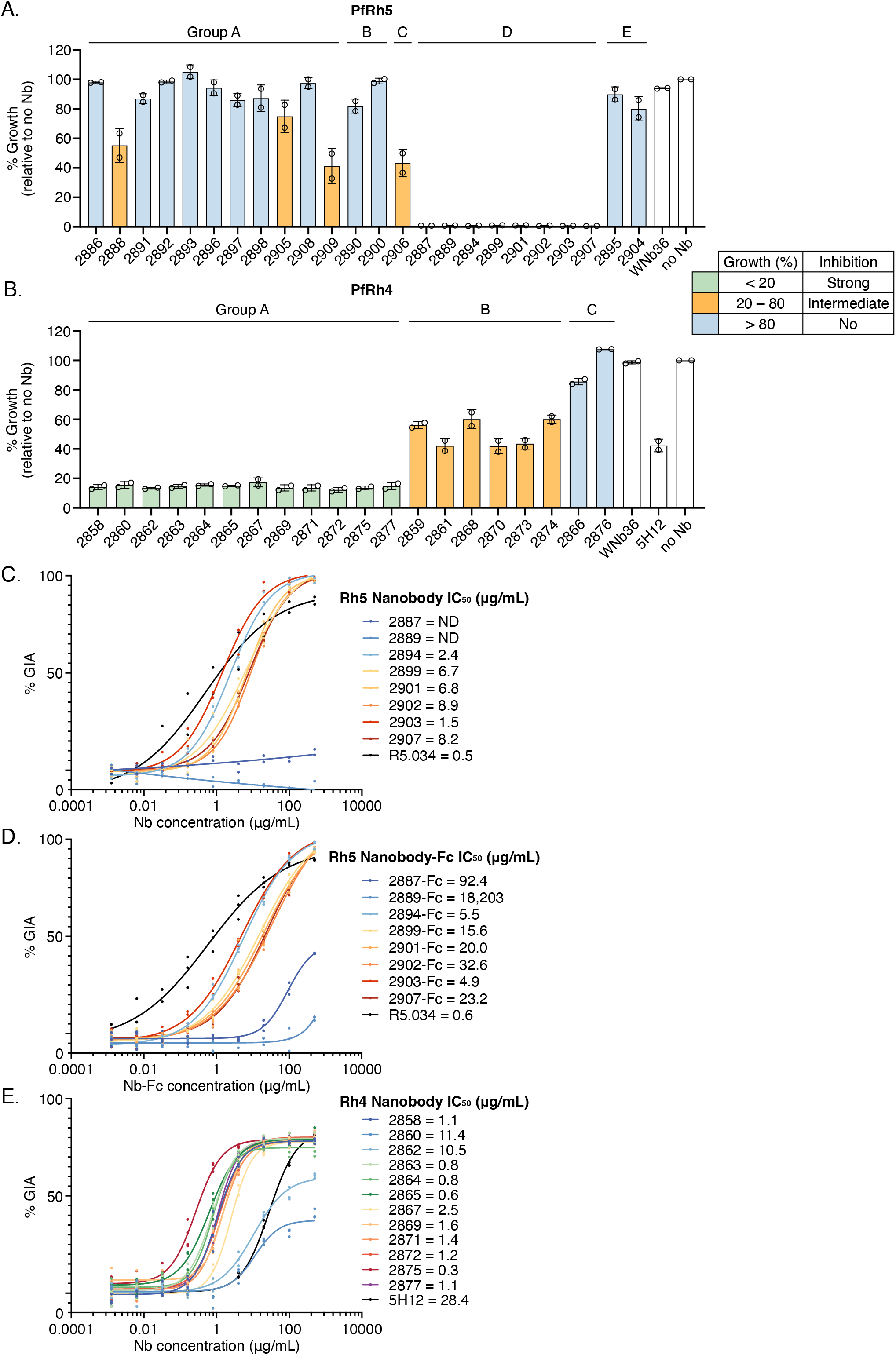
Nanobody inhibition of PfRh5 and PfRh4 invasion pathways. (A) Single concentration growth inhibition assays were performed using PfRh5 nanobodies at a final concentration of 0.5 mg/mL using the 3D7 strain of *P. falciparum*. (B) Single concentration growth inhibition assays were performed using PfRh4 nanobodies at a final concentration of 0.1 mg/mL with neuraminidase-treated erythrocytes using the D10-PHG strain of *P. falciparum*. Growth inhibitory activity IC_50_ values for (C) PfRh5 nanobodies, (D) PfRh5 nanobody-Fcs and (E) PfRh4 nanobodies were determined through a 9-point, 1:5 dilution series ranging from 500 – 0.00128 μg/mL. Resulting data were log transformed and fitted with a 4-parameter non-linear regression curve to calculate the IC_50_ value. Data for A and B are presented as mean ± SD, with each circle representing one independent experiment (n=3). Data for C, D and E are presented as individual points, representing the result for each concentration per replicate (for C, n = 2, for D and E, n = 3). ND not determined. Parasitaemia was measured after one replication cycle. Growth in the absence of nanobody was defined as 100%.

The 24 PfRh5 nanobodies were initially assessed for growth inhibitory activity at 0.5 mg/mL using the *P. falciparum* parasite strain 3D7 (Figure 3A). Eight nanobodies from Group D showed strong inhibition (Figure 3A). Four PfRh5 nanobodies showed intermediate inhibition, with three from Group A (2888, 2905 and 2909), and one Group C nanobody (2906). The remaining 12 PfRh5 nanobodies showed no growth inhibition (Figure 3A). As expected, the negative control SARS-CoV-2 nanobody WNb36, did not inhibit parasite invasion[29].

The 20 PfRh4 nanobodies were assessed at concentration of 0.1 mg/mL using neuraminidase-treated red blood cells as previously described[13,25,26,30], using the *P. falciparum* strain D10-PHG[31] (Figure 3B). We observed that all 12 of the PfRh4 Group A nanobodies were strongly inhibitory (Figure 3B). The six Group B nanobodies showed intermediate inhibitory activity and the two Group C nanobodies showed no inhibitory activity (Figure 3B). Negative control WNb36 showed no inhibition and the previously characterized mouse mAb 5H12 showed intermediate inhibition (Figure 3B).

We determined the IC_50_ values of the PfRh5 and PfRh4 inhibitory nanobodies. For the eight PfRh5 nanobodies that inhibited growth, we observed IC_50_ values ranging from 1.5 to 8.9 μg/mL and used monoclonal antibody R5.034 as a benchmarker [22] (Figure 3C, Data S3). In addition, we generated nanobody-Fcs and observed the IC_50_ values ranging from 4.9 to 18,203 μg/mL (Figure 3D, Data S3). These PfRh5 nanobody-Fcs bound to recombinant PfRh5 with nanomolar affinity with *K*_D_ values ranging from <0.01 nM to 6.71 nM (Data S1). In our assays, R5.034 showed the highest potency at 0.5 to 0.6 μg/mL compared to both PfRh5 monomeric nanobodies or nanobody-Fcs (Figure 3C and 3D). The 12 inhibitory PfRh4 nanobodies showed IC_50_ values ranging from 0.3 to 11.4 μg/mL (Figure 3E, Data S3).

### Computational design of stabilized PfRh4 variants encompassing the conserved structural scaffold

Members of the PfRh and PvRBP family share a conserved structural scaffold within their N-terminal domains, known to be important for human red blood cell receptor interactions [14,15,17,19]. PfRh5 represents the first crystal structure highlighting this structural scaffold among the *Plasmodium* PfRh and PvRBP families [17,18]. We showed that almost all PvRBP2b nanobodies bound to the conserved structural scaffold as well as the previously characterized stabilized PvRBP2b variant 2483[32] (Data S2, Figure 4A). We wanted to determine if this was also the case for PfRh4 nanobodies. However, to date, we have been unsuccessful in generating this conserved structural scaffold domain in PfRh4 with reliable yields (Figure 4Ai). Using computational approaches, we designed four PfRh4 variants encompassing amino acids 102 to 442 (2739, 2740, 2741 and 2742, referred to as 39, 40, 41 and 42) (Figure 4A). Each variant contained between 14 - 49 mutations distributed across both the protein core and surface residues (Table S1). For each of these, a corresponding paired variant was designed with minimal surface mutations, incorporating 10 to 28 mutations within the protein core (2743, 2744, 2745 and 2746, referred to as 43, 44, 45 and 46) (Table S1).

**Figure 4.**
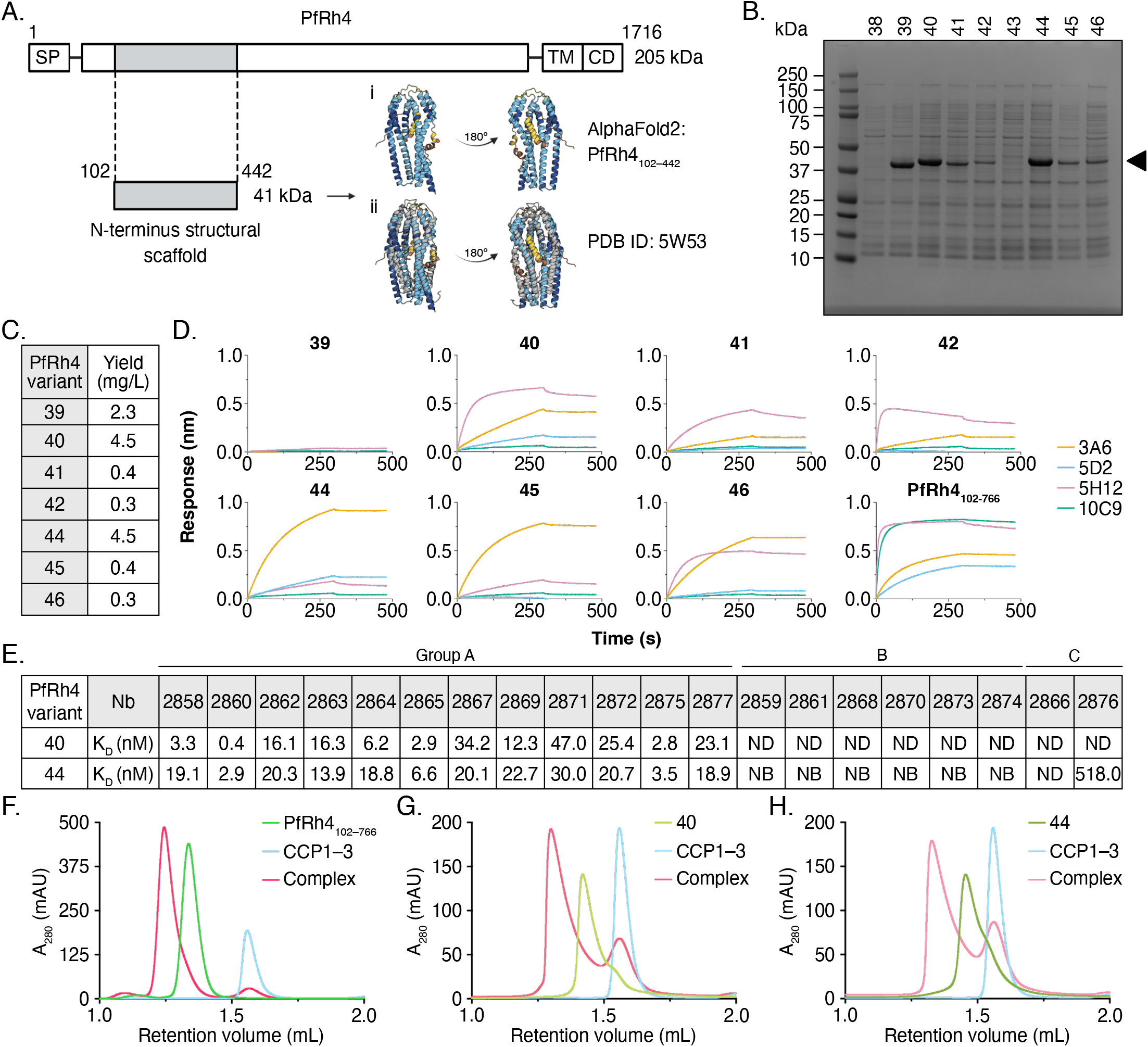
Expression of stabilized PfRh4 variants. (A) Schematic of wild-type PfRh4_1–1716_ and its N-terminal structural scaffold, aa 102–442 used for variant design with (i) AlphaFold2 predicted structure of PfRh4_102–442_. Structure colored based on model confidence where dark blue is high confidence and orange is very low confidence. (ii) Superimposition of the N-terminal domain crystal structure of PvRBP2b (grey, PDB ID: 5W53) with AlphaFold2 prediction of PfRh4_102–442_. SP, signal peptide; TM, transmembrane domain; CD, cytoplasmic domain. (B) PfRh4 variants separated by SDS-PAGE gel under non-reducing conditions after Ni-NTA affinity chromatography (arrow indicated band ∼44 kDa). (C) Comparison of yields between variants after two to three step purification process. (D) Single binding analysis of PfRh4_102–766_ and variants to mouse monoclonal antibodies by BLI. Representative curves of two independent experiments are plotted. (E) Binding affinities (*K*_D_) of PfRh4 nanobodies for variants 40 and 44 determined by BLI. Values represent means from two independent experiments. NB, no binding; ND, not determined. (F) Analytical SEC of PfRh4 variants with CCP1–3. PfRh4_102–766_ is 80 kDa and elutes at 1.3 mL (lime green) and CCP1–3 is 21 kDa and elutes at 1.6 mL (light blue). The PfRh4_102–766_-CCP1–3 complex is 101 kDa and elutes at 1.2 mL (deep pink). (G) Analytical SEC of 40 with CCP1–3. Variant 40 is 40 kDa and elutes at 1.4 mL (light green). The 40-CCP1–3 complex is 61 kDa and elutes at 1.3 mL (pink). (H) Analytical SEC of variant 44 with CCP1–3. Variant 44 is 40 kDa and elutes at 1.4 mL (dark green). The 44-CCP1–3 complex is 61 kDa and elutes at 1.3 mL (light pink).

We expressed parental PfRh4 and the eight variants in *Escherichia coli* in small scale and used Ni-NTA affinity chromatography as the first purification step. Parental PfRh4 (2738, referred to as 38) and the eight variants are predicted to migrate at 44 kDa. SDS-PAGE analyses of purified proteins after Ni-NTA affinity chromatography showed that seven of the eight PfRh4 variants (39, 40, 41, 42, 44, 45 and 46) were expressed successfully (Figure 4B). Parental PfRh4 (38) and variant 43 showed little to no protein expression and were excluded from further expression and analysis (Figure 4B). Using 2 L *E. coli* cultures, we re-expressed the seven PfRh4 variants and purified them using a combination of Ni-NTA affinity chromatography, cation-exchange chromatography and size exclusion chromatography (SEC) if yields were permissible (Figure S3). PfRh4 variants 40 and 44 showed the highest expression levels at 4.5 mg per L of bacterial culture, variant 39 showed intermediate expression at 2.3 mg per L, while the remaining four variants had low yields ranging from 0.3 - 0.4 mg per L of *E. coli* culture (Figure 4C).

To assess whether the seven PfRh4 variants retained recognition of previously characterized PfRh4-specific mouse monoclonal antibodies (mAbs) [26], we performed single binding curve analysis using BLI (Figure 4D, Data S2). Recombinant PfRh4 spanning residues 102 to 766 was used as a positive control antigen, as it is known to bind all four PfRh4-specific mouse mAbs (3A6, 5D2, 5H12 and 10C9)[26]. The 3A6 and 5D2 mAbs bind to peptides spanning residues 238 to 280 of PfRh4, whereas 10C9 mAb recognizes a C-terminal epitope of PfRh4 spanning residues 662 to 680 and was used as a negative control mAb[26], as the stabilized PfRh4 variants only encompasses residues 102 to 442. A BLI response threshold of 0.1 nm was used to define positive binding.

PfRh4 variants 40 and 44 retained binding to all the three N-terminal specific mAbs 3A6, 5D2 and 5H12 (Figure 4D). Four variants, 41, 42, 45 and 46 retained binding to only 3A6 and 5H12 (Figure 4D). Variant 39 showed no detectable binding to any of the three mAbs (Figure 4D). As expected, no binding was observed to 10C9 for any of the seven PfRh4 variants.

### PfRh4 variants 40 and 44 bind PfRh4 nanobodies and retain binding to human receptor Complement Receptor 1

Both PfRh4 variants 40 and 44 also bound all Group A nanobodies with *K*_D_ values ranging from 0.4 to 47.0 nM (Figure 4E) but showed no measurable binding to nanobodies from Group B and Group C (Figure 4E). The only exception was variant 44, which retained binding to nanobody 2876 with a *K*_D_ of 518.0 nM (Figure 4E). To evaluate whether PfRh4 variants 40 and 44 retained the ability to interact with a fragment of CR1, CCP1–3, we performed analytical SEC (Figure 4F - H). The interaction between PfRh4_102–766_ and CCP1–3 was used as a positive control. We ran CCP1–3 (light blue, Figure 4F) and PfRh4_102–766_ (lime green, Figure 4F) individually. Incubation of CCP1–3 and PfRh4_102–766_ together resulted in complex formation (deep pink, Figure 4F). Incubation of 40 and 44 with CCP1-3 resulted in complex formation, which shows that this smaller fragment of PfRh4 from residues 102-442 retains the ability to engage with CR1 (deep pink, Figure 4G and 4H respectively).

## Discussion

To the best of our knowledge, we present the first collection of nanobodies targeting key malaria parasite invasion adhesins; PfRh4, PfRh5 and PvRBP2b. Nanobodies specific to PfRh5, PfRh4 and PvRBP2b were identified that bind their respective targets with high affinity in the nanomolar range. This collection comprises nanobodies that bind three to eight distinct epitopes across PfRh5, PfRh4 and PvRBP2b. Several of these nanobodies effectively block receptor binding and/or inhibit parasite invasion of red blood cells. We also employed structure-guided protein design to successfully express stabilized variants of the N-terminal domain of PfRh4 spanning the conserved structural scaffold present in the PfRh and PvRBP families, which retained mAb, nanobody and CR1 receptor recognition.

All PfRh5 nanobodies that blocked complex formation with basigin also inhibited parasite invasion, suggesting that these nanobodies likely target the basigin binding site on PfRh5. The most potent PfRh5 human monoclonal antibody R5.034 inhibits parasite invasion through a mechanism that does not directly block the interaction between PfRh5 and basigin [22], suggesting that our nanobodies use a distinct inhibitory mechanism compared to R5.034. We show that R5.034 was still the most potent antibody in comparison to our monomeric nanobodies or nanobody-Fcs. Structural characterization of PfRh5-nanobody complexes in the future will be essential to define the precise epitopes targeted by the nanobodies which enable them to block receptor engagement and inhibit parasite invasion. In addition, all PfRh4 nanobodies that blocked complex formation with CR1 also inhibited parasite invasion of neuraminidase-treated red blood cells. The 12 PfRh4 nanobodies showed 2 to 10-fold higher potency compared to 5H12, a mouse monoclonal antibody [26].

Crystal structures of the N-terminal domain of PfRh5, PvRBP2b and PvRBP2a highlight the conserved structural scaffold that unites the PfRh and PvRBP family of adhesins [14,15,17,19]. In previous studies, we have not been able to express the N-terminal domain of PfRh4 (residues 102–442) in bacterial expression systems to sufficient yields. Using computational protein design, we successfully generated stabilized variants of the N-terminal domain of PfRh4 that expressed at yields of up to 4.5 mg per L in bacterial culture. Among these, two of the top expressing PfRh4 variants retained their ability to bind both PfRh4 mouse monoclonal antibodies and nanobodies, as well as the PfRh4 receptor, CCP1–3 of CR1.

One limitation of this study is that the engineered PfRh4 variants exhibited reduced affinity to Group A nanobodies and loss of recognition to Group B and C nanobodies. This suggests that while the expression of the PfRh4_102-442_ domain was improved, the design process introduced mutations that were deleterious for high affinity nanobody recognition of certain epitopes. However, the loss of binding to Group B and C nanobodies may be due to the epitopes of these nanobodies residing within residues 443-766, which is present in the immunization antigen PfRh4_102-766_, but not in the PfRh4 variants spanning residues 102-442. To improve our protein engineering designs to maintain enhanced expression and the preservation of native epitope integrity, we will require high resolution crystal structures of these variants in future work.

## Materials and Methods

### Alpaca immunization and isolation of PfRh4, PfRh5 and PvRBP2b nanobodies

PfRh5 (residues 127–526), PfRh4 (residues 102–766) and PvRBP2b (residues 161–1454) were used to immunize individual alpacas. Alpacas were immunized six times with approximately 200 μg of recombinant antigen on days 0, 14, 21, 28, 35 and 42. The adjuvant used was GERBU FAMA. Immunization and handling of the alpacas for scientific purposes was approved by Agriculture Victoria, Wildlife & Small Institutions Animal Ethics Committee, project approval No. 26-17. Blood was collected three days after the last immunization for the preparation of lymphocytes. Nanobody library construction was carried out according to established methods as described[33]. Briefly, alpaca lymphocyte mRNA was extracted and amplified by RT-PCR with specific primers to generate a cDNA nanobody library. The library was cloned into a pMES4 phagemid vector and amplified in *E. coli* TG1 strain and subsequently infected with M13K07 helper phage for recombinant phage expression. Biopanning for PfRh5, PfRh4 and PvRBP2b nanobodies using phage display was performed as previously described with the following modifications[34–37]. Phage displaying antigen-specific nanobodies were enriched after two rounds of biopanning with 1 μg of antigen immobilized on Dynabeads M-280 streptavidin (Thermo Fisher). After the second round of panning, phage supernatants were screened for positive hits using ELISA.

### Nanobody phage ELISA

96-well flat-bottom MaxiSorp plates (Nunc) were coated with 62 nM of antigen in 50 μL of PBS for 1 hour at room temperature. All washes were done three times using PBS containing 0.05% (v/v) Tween-20 (PBST) and all incubations were performed for 1 hour at room temperature. Antigen coated plates were washed and blocked by incubation with 10% (w/v) skim milk in PBST. Overnight cultures of individual phage clones grown in 96 deep well plates were centrifuged at 600 *x* g for 10 min at 4°C to pellet bacteria. After blocking, ELISA plates were washed and incubated with 50 µL of phage supernatant. The plates were washed and incubated with 50 µL of horseradish peroxidase (HRP)-conjugated anti-M13 (Abcam; 1:15,000) diluted in 2% (w/v) skim milk in PBST. Following the final wash, 50 μL of azino-bis-3-ethylbenthiazoline-6-sulfonic acid liquid substrate (ABTS; Sigma) was added and incubated at room temperature. The reaction was stopped with 50 μL of 1% (v/v) SDS and absorbance was measured at 405 nm. All positive clones were sequenced using Sanger sequencing and annotated using PipeBio. Nanobody clonal groups were defined by less than three amino acid differences in the CDR3.

### Next-Generation Sequencing and analyses of nanobody phage libraries

Plasmids from phage libraries were digested and gel purified to isolate the nanobody domain sequences. Paired-end 2×300 bp sequencing libraries were prepared using the NEBNext Multiplex Oligos for Illumina (Cat # E7395) in a PCR-free manner according to the manufacturer’s instructions. The samples were size-selected using AMPure XP beads, quantified by qPCR using the KAPA Library Quantification Kit and sequenced on an Illumina NextSeq 2000 instrument. Raw data processing and most analysis was performed using alpseq v0.5.0 [28] with default settings. Cladograms were generated from generalized Levenshtein distances between CDR3s with the Neighbor-Joining method[38] from the phangorn (v2.12.1) [39] package and plotted using the ggtree (v3.14.0) [40] package in R (v4.4.1) [41].

### Nanobody expression and purification

Nanobodies were expressed in *E. coli* WK6 cells. Bacteria were grown in Terrific Broth supplemented with 0.1 % (w/v) glucose and 100 µg/mL carbenicillin at 37°C at 180 rpm to an OD_600_ of 0.7 and induced with 1 mM IPTG and grown overnight at 28 °C for approximately 16 hours. For periplasmic extraction of nanobodies, cell pellets were harvested and resuspended in 20 % (w/v) sucrose, 10 mM imidazole pH 7.5, 150 mM NaCl, DPBS and incubated for 15 min on ice. EDTA pH 8.0 was then added to a final concentration of 5 mM and incubated on ice for 20 min. To prevent EDTA chelation of the nickel columns, 10 mM MgCl_2_ was added, and periplasmic extracts were harvested by centrifugation at 30,000 *x* g for 30 mins. The supernatant was filtered using a 0.22 µm filter and loaded onto a 1 ml HisTrap Excel HP column (Cytiva Cat# 17371205) which was pre-equilibrated in 5 mM imidazole pH 7.5, 100 mM NaCl, DPBS. Nanobodies were eluted using 400 mM imidazole pH 7.5, 100 mM NaCl, DPBS and concentrated. They were buffer exchanged into sterile DPBS using 3K MWCO Amicon Ultra-15 centrifugal filter unit concentrators (Merck Millipore).

### Biolayer interferometry for affinities

Nanobody affinity measurements were performed using the Octet RED96e instrument (FortéBio). Assays were performed at 25°C in solid black 96-well F-bottom plates (Greiner-Bio One #655209) agitated at 1,000 rpm. Kinetic buffer consisted of PBS with 0.1% (w/v) BSA and 0.05% (v/v) Tween-20. A 60 s baseline step was applied before nanobodies were loaded onto His1K biosensors (Sartorius) by submerging sensor tips in 5 µg/mL of each nanobody until a response of 0.5 nm was obtained. Sensors were then dipped in kinetic buffer for 60 s. Association measurements were performed for 180 s using a two-fold serial dilution of PfRh5 from 6 to 100 nM and dissociation was measured in kinetic buffer for 180 s. Sensors were regenerated using a cycle of 5 s in 100 mM glycine pH 1.5 and 5 s in kinetic buffer repeated five times. Baseline drift was corrected by subtracting the response of a nanobody-loaded sensor dipped in kinetic buffer for the association step. Curve fitting analysis was performed with Octet Data Analysis 11.1 software using a global fit 1:1 model to determine *K*_D_ values and kinetic parameters. Curves that could not be fitted well with R^2^ >0.98 and X^2^ <0.5 were excluded from the analyses.

Nanobody affinities to PfRh4 and PvRBP2b were measured with the above method, using a two-fold serial dilution of PfRh4 from 6 to 100 nM and PvRBP2b from 6 to 200 nM for higher affinity nanobodies and 100 to 3200 nM for lower affinity nanobodies. Nanobody affinities to PfRh4 variants 2740 and 2744 were measured with the above method, using a two-fold serial dilution from 13 to 400 nM for higher affinity nanobodies and 250 to 2000 nM for lower affinity nanobodies.

### Biolayer interferometry for epitope binning

For epitope binning experiments, 15 nM PfRh5 was pre-incubated with each nanobody at a 10-fold molar excess (150 nM) for 1 hour at room temperature. A 30 s baseline step was established between each step of the assay. His1K sensors were first loaded with 10 µg/mL of nanobody for 5 min. The sensor surface was quenched by dipping into 10 µg/mL of an irrelevant nanobody (WNb27 against SARS-CoV-2) for 5 min. Nanobody-loaded sensors were then dipped into premixed solutions of PfRh5 and nanobody for 5 min. Nanobody loaded sensors were also dipped into PfRh5 alone to determine the level of PfRh5 binding to immobilized nanobody in the absence of other nanobodies. Percentage competition was calculated by dividing the maximum response of the premixed PfRh5 and nanobody solution binding by the maximum response of PfRh5 binding alone, multiplied by 100. Competition was defined as < 40% binding. Epitope binning experiments with PfRh4 and PvRBP2b were performed using the above method with the following modification. 10 nM PfRh4 was pre-incubated with each nanobody at a 10-fold molar excess (100 nM), and 500 nM of PvRBP2b was pre-incubated with each nanobody at a 10-fold molar excess (5000 nM).

### Fluorescence resonance energy transfer (FRET) assay

Basigin and PfRh5 were labelled with *N*-hydroxysuccinimide ester-activated DyLight 488 (DL488) and DyLight 594 (DL594) (Life Technologies) respectively. The dyes were dissolved in dimethyl sulfoxide (DMSO) (Sigma) and added at three-fold molar excess to protein. After 1 hour incubation at room temperature, unconjugated dye was removed by buffer exchange into 20 mM HEPES pH 7.5, 150 mM NaCl using 3K MWCO Amicon Ultra-4 centrifugal filter unit concentrators (Merck Millipore) at 4000 *x* g. Average dye per protein was ∼1.4 dye/protein for basigin and ∼2.1 dye/protein for PfRh5. Basigin-DL488 and PfRh5-DL594 were mixed in a 1:1 molar ratio with 10-fold excess of nanobodies in a final reaction volume of 10 μL in FRET buffer (20 mM HEPES pH 7.5, 150 mM NaCl and 0.01% (v/v) BSA) and read in Corning 384-well plates. Each sample was performed in triplicate. 1 μL of 2% (v/v) SDS was added to one triplicate sample with no nanobodies to measure the background signal. Fluorescence intensity was measured using CLARIOstar Plus (BMG Labtech). DL488 fluorescence was measured with a 488/14-nm excitation filter and 535/30-nm emission filter and DL594 fluorescence was measured with a 575/20-nm excitation filter and 630/40-nm emission filter. Sensitized emission was measured with a 488/14-nm excitation filter and 630/40-nm emission filter. Fluorescence measures were analyzed using Prism software (GraphPad; version 10.4.1). To account for bleed-through of dyes into the sensitized emission spectra, standard curves for both dyes were measured and the raw data was transformed by multiplying with the slope of the standards. The *y*-intercept from the standard curves indicated the background fluorescence when no dye was present. The FRET signal was calculated with the following equation: FRET signal = Raw sensitized emission - transformed DL488 emission - transformed DL594 emission - DL488 standard curve *y*-intercept - DL594 standard curve *y*-intercept. To compare between independent replicates, the FRET signal was normalised by dividing the FRET signal with nanobody by the FRET signal with no nanobody, multiplied by 100.

FRET with PfRh4 specific nanobodies was performed using the above method. CCP1-2 of CR1 and PfRh4 were labelled with DL488 and DL594, respectively. Average dye per protein was ∼1.1 dye/protein for CCP1-2 and ∼2.3 dye/protein for PfRh4. FRET buffer used was 50 mM MES pH 6.0, 150 mM NaCl, 0.01% (v/v) BSA. PvRBP2b and TfR1 were labelled with DL488 and DL594, respectively. Average dye per protein was ∼1.4 dye/protein for PvRBP2b and ∼2.6 dye/protein for TfR1. FRET buffer used was 20 mM HEPES pH 7.5, 150 mM NaCl, 0.01% BSA.

### Growth Inhibition Assay

Growth inhibition assays were performed using either *P. falciparum* 3D7 or D10-PHG strains. Parasite culture was maintained using O^+^ red blood cells (Australian Red Cross, Lifeblood) at 4 % haematocrit in RPMI 1640, GlutaMAX, HEPES (Thermo Fisher) supplemented with: hypoxanthine (12.5 mg/L) (Sigma), D-glucose (2 g/L) (Thermo Fisher), gentamycin (20 mg/L) (Thermo Fisher), 0.5 % (w/v) AlbuMAX II (Thermo Fisher). Assays were set up using trophozoite stage parasites at 1 % parasitaemia, 2 % haematocrit in 96-well U bottom plates. Assay volumes were 50 μL per well, including 5 uL of relevant nanobody or control treatment. Nanobodies were assessed at 0.5 mg/mL or 0.1 mg/mL for single concentration assays, or 500 – 0.00128 µg/mL for IC_50_ titration assays. Plates were incubated at 37ºC, in 1% O_2,_ 5% CO_2_, balance N_2_ for 48 hours to allow a single replication cycle to proceed. All wells were then stained with 1X SYBR Green I (Invitrogen) diluted in PBS, and parasitaemia was determined using the Novocyte Quanteon Flow Cytometer (Agilent). For each well, a total of 90,000 red blood events were collected (gated on SSC-A vs. FSC-A, then again on FSC-H vs FSC-A). Parasitised cells were gated using the B530 detector (FITC) vs. FSC-A. Parasite growth was calculated with the following equation: Growth (% PBS/no nanobody) = (parasitaemia % treated/parasitaemia % control)*100. Growth inhibitory activity was calculated with the following equation: % GIA = 100 – ([parasitaemia % treated/parasitaemia % control] *100). Samples were tested in technical duplicates.

### PROSS stability design

Due to the limited sequence diversity among natural PfRh4 homologs, we employed a hybrid computational design strategy combining the AI-based inverse-folding algorithm ProteinMPNN and the PROSS workflow[32,42] which is based on Rosetta atomistic calculations and an analysis of the natural sequence diversity of homologs. As the molecular structure of PfRh4_102–442_ has not been determined experimentally, we used AlphaFold2 to generate a predicted structure. This model was used to generate a backbone-specific mutation model by ProteinMPNN[43]. Log-likelihoods for amino acid substitutions at each amino acid position were extracted from the model and averaged over 50 independent model initialisations, then rounded to the nearest integer to account for stochastic variations. The resulting log-probability matrix was used as replacement of the position-specific scoring matrix (PSSM) that is used by PROSS to assess the natural sequence diversity of homologs. Structural design calculations followed the standard PROSS protocol as the energy function, a PSSM score threshold ≥ −2 and positions with pLDDT >= 50 to determine designable positions. Mutations at positions deemed too risky were manually discarded, leaving only four designable positions in low-confidence segments (50 <= pLDDT <= 70): positions 138, 270, 274, and 276.

### Expression and purification of parental PfRh4 and variants

PfRh4 sequence from *P. falciparum* 3D7 was obtained from PlasmoDB Database (www.plasmodb.org; accession number: PF3D7_0424200, 1716 amino acids). Synthetic DNA was codon-optimised for expression in *E. coli*. The DNA fragment encoding amino acids 102 to 442 of PfRh4, including an N-terminal 6xHis-tag and TEV protease cleavage site was cloned using restriction enzyme-based cloning into pET-45 vector. This sequence refers to the parental PfRh4 or variant 38. Restriction enzyme cloning was used to clone the synthetic DNA fragments (obtained from Twist Biosciences) of PfRh4_102–442_ variants into the same pET-45 vector. All positive plasmids were sequence verified at the WEHI Advanced Genomics Facility.

Parental PfRh4 (38) was expressed using *E. coli* SHuffle T7 Express cells and Terrific Broth supplemented with 100 μg/ml of carbenicillin. Flasks containing 1 L of medium were incubated in a Multitron shaker (Infors HT) at 37°C at 180 rpm. At OD_600_ of approximately 0.8, IPTG was added to the final concentration of 1.0 mM and grown overnight for 18 h at 16°C. Cells were harvested by centrifugation at 6000 *x g* and resuspended in freezing buffer containing 50 mM Tris-HCl pH 7.5, 500 mM NaCl, 10 % (v/v) glycerol supplemented with cOmplete EDTA-free protease inhibitor cocktail, and stored at −80°C until further processing.

For purification, cell pellet was thawed on ice and resuspended in the freezing buffer containing 0.5 mg/ml of DNase, 1.0 mg/ml of lysozyme and MgCl_2_ to a final concentration of 5mM. Cells were lysed by sonication using the Sonopuls UW 3200 equipped with VS 70 T probe. The lysate was clarified by centrifugation at 30,000 *x g* for 45 min at 4°C. The supernatant was loaded onto a 5 ml HisTrap Excel column pre-equilibrated with the freezing buffer. Unbound material was removed using 10 column volumes of Wash Buffer: 20 mM Tris–HCl pH 7.5, 500 mM NaCl and 10 mM imidazole pH 7.5. The bound protein was eluted from the column using the Elution buffer (Wash Buffer supplemented with 300 mM imidazole pH 7.5). Eluted fractions were pooled, and TEV protease was added before being dialyzed overnight into the dialysis buffer containing 50 mM MES pH 6.0 and 150 mM NaCl. The resulting protein sample was applied on the 5 ml HiTrap SP HP cation exchange chromatography column pre-equilibrated with the dialysis buffer. Unbound material was removed using 10 column volumes of the buffer. Protein was eluted from the column using a salt gradient of 0 to 1.0 M NaCl in 50 mM MES pH 6.0. Collected fractions were analyzed on SDS-PAGE and fractions of interest were concentrated using an Amicon Ultra-4 10 kDa molecular weight cut-off concentrator and loaded onto HiLoad Superdex 75 16/600 size-exclusion column pre-equilibrated with 50 mM MES pH 6.0 and 150 mM NaCl. The monodisperse peak fractions containing protein were pooled and concentrated using the same concentrator, flash-frozen in liquid nitrogen and stored at −80°C. Expression and purification of PfRh4 variants 39 – 46 were performed in a similar manner as described above.

### Analytical SEC for complex formation

Analytical SEC was performed on the ÄKTA pure 25 M1 chromatographic system using a Superdex 200 Increase 3.2/300 column pre-equilibrated with 50 mM MES pH 6.0 and 150 mM NaCl. PfRh4_102–766_ was first injected onto the column by itself at 50 µM in 50 µL to create a reference point for individual proteins followed by CCP1–3. PfRh4_102–766_ was then mixed with CCP1–3 at a 1:1 molar ratio and incubated for 1 hour at room temperature in a final volume of 50 µL before being injected onto the column. Elution was monitored with the absorbance signal at 280 nm and fractions were collected to analyze on an SDS-PAGE gel. Analytical SEC with PfRh4 variants 40 and 44 were performed using the above method. Expression and purification of CR1 fragments were as previously described [26].

## Data Availability Statement

All data is available in the manuscript and supplementary materials. Nanobody sequences used in the manuscript is available in Supplementary Dataset 1.

## Acknowledgments

We thank Stephen Scally for providing us with recombinant basigin. W-H.T. is supported by National Health and Medical Research Council of Australia (NHMRC) GNT2016908 and APP2001385. Q.G. is supported by NHMRC Investigator Grant GTN2007996. We thank the Australian Red Cross Lifeblood for the supply of human red blood cells and serum. The authors acknowledge the Victorian State Government Operational Infrastructure Support and Australian Government NHMRC IRIISS.

## Supplementary data

**Figure S1. Sanger sequencing summary and epitope binning of PfRh5, PfRh4 and PvRBP2b specific nanobodies.** The nanobody clonal groups from phage display panning against PfRh5 (A), PfRh4 (C) and PvRBP2b (E) identified using Sanger sequencing. The number in the inner circle indicates the number of distinct clonal groups based on the CDR3 sequences. Colored pie slices are proportional to the number of clonally related sequences. Epitope binning competition experiments using BLI with PfRh5 (B), PfRh4 (D) and PvRBP2b (F). Immobilized nanobodies are indicated on the left columns and nanobodies pre-incubated with antigens at a 10:1 molar ratio are indicated on the top rows. Binding of antigen pre-mixed with nanobody was calculated relative to antigen binding alone, which was assigned to 100%. The green and white boxes represent non-competing and competing nanobodies, respectively.

**Figure S2. Selection hotspots within the sequence space of observed nanobody variation.** (A) Each point represents a unique nanobody sequence that has been embedded into a vector using the AntiBERTy antibody language model and reduced into 2D space for visualisation via UMAP. Grey represents all observed nanobody variation prior to phage display selection. Nanobody sequences remaining after the second round of phage display selection against PfRh5, PfRh4 and PvRBP2b respectively are coloured according to their abundance (log10 of CPM). (B) Sequences highlighted in orange are from the same clonal group as nanobodies selected for characterisation in this study against PfRh5, PfRh4 and PvRBP2b.

**Figure S3. Purification of stabilized PfRh4 variants.** (A) Schematic of the three-step purification process for PfRh4 variants. Variants taken through each purification step are listed below each step. (B) SEC chromatogram and corresponding reduced SDS-PAGE gels of variants. Fractions that were pooled as the final purified protein are indicated by the dotted lines and corresponding box on the SDS-PAGE gel. I, input. Created in https://BioRender.com

**Table S1. Mutated residues between PfRh4**_**102–442**_ **and variants**.

**Data S1. Data relating to nanobody characterization performed in Figure 1.** Table of nanobody CDR3 sequences, full-length (FL) sequences, their corresponding counts per million (CPM) from NGS, enrichment from Sanger sequencing and determined binding kinetics and affinity data. Data are from two independent biolayer interferometry experiments and shown are the mean ± SD values for affinity (*K*_D_), association rate (*k*_a_) and dissociation rate (*k*_d_). Chi-squared (X^2^) and R-squared (R^2^) are statistical values indicating quality of fit between experimental data and theoretical model. ND, not determined; NB, no binding; NE, no expression.

**Data S2. Single binding data for PvRBP2b and PfRh4 nanobodies against the N-terminal conserved scaffold**. (A) Binding response of PvRBP2b nanobodies to PvRBP2b_161–1454_, PvRBP2b_169–470_ and PvRBP2b_169–470_ stabilized design (2483) from one experiment as measured by biolayer interferometry. (B) Binding response of PfRh4_102–766_ mouse monoclonal antibodies to PfRh4_102–766_ and variants from two independent experiments with mean ± SD values shown.

**Data S3**. Replicate IC_50_ values for PfRh5 nanobodies, PfRh5 nanobody-Fcs and PfRh4 nanobodies.

