## Supplemental Figures and Tables for "Nanobodies against *Plasmodium* adhesins that block receptor engagement and malaria parasite invasion"

Figure S1

A.

PfRh5

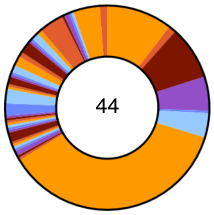


B.

**Competing nanobody**

| Immobilised nanobody | 2886 | 2888 | 2891 | 2892 | 2893 | 2896 | 2897 | 2898 | 2905 | 2908 | 2909 | 2890 | 2900 | 2906 | 2887 | 2889 | 2894 | 2899 | 2901 | 2902 | 2903 | 2907 | 2895 | 2904 | Group |
|---|---|---|---|---|---|---|---|---|---|---|---|---|---|---|---|---|---|---|---|---|---|---|---|---|---|
| 2886 | 0 | 2 | 1 | 1 | 5 | 1 | 3 | 1 | 1 | 5 | 6 | 3 | 3 | 106 | 89 | 86 | 85 | 90 | 88 | 92 | 84 | 91 | 108 | 105 | Group A |
| 2888 | 3 | 7 | 4 | 2 | 7 | 3 | 5 | 2 | 1 | 6 | 5 | 4 | 3 | 101 | 92 | 85 | 85 | 88 | 87 | 92 | 83 | 89 | 110 | 105 | |
| 2891 | 2 | 5 | 3 | 2 | 8 | 3 | 9 | 2 | 1 | 5 | 5 | 4 | 4 | 101 | 90 | 85 | 86 | 88 | 86 | 91 | 81 | 89 | 107 | 104 | |
| 2892 | 2 | 2 | 3 | 2 | 6 | 2 | 3 | 1 | 2 | 4 | 5 | 3 | 3 | 100 | 87 | 85 | 81 | 86 | 86 | 91 | 81 | 88 | 107 | 104 | |
| 2893 | 1 | 2 | 2 | 1 | 4 | -0 | 2 | 0 | 2 | 2 | 5 | 2 | 2 | 94 | 91 | 88 | 87 | 91 | 92 | 96 | 88 | 89 | 110 | 108 | |
| 2896 | 2 | 2 | 2 | 2 | 4 | 1 | 3 | 1 | 0 | 3 | 4 | 2 | -0 | 88 | 89 | 87 | 84 | 86 | 90 | 94 | 84 | 91 | 110 | 106 | |
| 2897 | 2 | 5 | 3 | 1 | 5 | 2 | 4 | 1 | 2 | 3 | 5 | 3 | 1 | 94 | 90 | 84 | 83 | 83 | 86 | 92 | 82 | 86 | 108 | 104 | |
| 2898 | 1 | 5 | 3 | 1 | 7 | 4 | 13 | -0 | 1 | 4 | 6 | 3 | 3 | 105 | 92 | 89 | 89 | 91 | 90 | 94 | 86 | 92 | 106 | 103 | |
| 2905 | 1 | 5 | 4 | 1 | 8 | 3 | 8 | 2 | 1 | 4 | 6 | 3 | 3 | 105 | 91 | 86 | 87 | 86 | 87 | 92 | 83 | 88 | 107 | 105 | |
| 2908 | -0 | 0 | -0 | 1 | 6 | -1 | 1 | -0 | 0 | 2 | 4 | 1 | -1 | 87 | 85 | 83 | 85 | 86 | 88 | 93 | 84 | 91 | 109 | 107 | |
| 2909 | 0 | 1 | 2 | 3 | 7 | 0 | 2 | 1 | 1 | 6 | 5 | 2 | 1 | 96 | 85 | 83 | 82 | 82 | 85 | 91 | 82 | 87 | 107 | 105 | |
| 2890 | 1 | 4 | 3 | 1 | 6 | 2 | 4 | 2 | 2 | 4 | 5 | 3 | 1 | 78 | 89 | 84 | 82 | 86 | 86 | 91 | 82 | 88 | 106 | 103 | Group B |
| 2900 | 1 | 2 | 2 | 2 | 5 | 1 | 3 | 2 | 2 | 3 | 6 | 2 | 2 | 62 | 90 | 87 | 84 | 90 | 90 | 94 | 87 | 87 | 109 | 106 | |
| 2906 | 79 | 83 | 76 | 66 | 94 | 56 | 72 | 112 | 53 | 46 | 82 | 21 | 13 | 7 | 90 | 89 | 84 | 86 | 88 | 94 | 84 | 90 | 113 | 107 | Group C |
| 2887 | 109 | 105 | 108 | 109 | 110 | 102 | 105 | 129 | 101 | 109 | 103 | 113 | 111 | 118 | 1 | 3 | 1 | 1 | 2 | 0 | -0 | 3 | 127 | 116 | Group D |
| 2889 | 108 | 104 | 106 | 108 | 110 | 100 | 102 | 124 | 99 | 109 | 103 | 111 | 107 | 114 | 1 | 1 | 1 | -1 | 1 | 1 | 1 | 1 | 128 | 117 | |
| 2894 | 108 | 105 | 108 | 108 | 109 | 104 | 107 | 122 | 102 | 107 | 104 | 111 | 109 | 115 | 1 | 3 | 1 | 1 | 2 | 1 | -0 | 2 | 123 | 115 | |
| 2899 | 109 | 106 | 105 | 109 | 111 | 101 | 103 | 126 | 101 | 107 | 103 | 111 | 109 | 116 | 3 | 4 | 3 | 3 | 2 | 2 | 1 | 2 | 123 | 117 | |
| 2901 | 108 | 104 | 107 | 108 | 110 | 103 | 105 | 126 | 100 | 106 | 101 | 112 | 109 | 114 | 1 | 3 | 1 | 1 | 1 | 0 | -1 | 2 | 127 | 117 | |
| 2902 | 109 | 111 | 107 | 109 | 113 | 101 | 103 | 129 | 101 | 108 | 105 | 112 | 111 | 116 | 6 | 3 | 4 | 3 | 2 | 4 | 4 | 3 | 129 | 118 | |
| 2903 | 109 | 106 | 107 | 110 | 111 | 103 | 105 | 123 | 102 | 109 | 105 | 111 | 108 | 115 | 2 | 2 | 1 | -0 | 2 | 2 | 1 | 1 | 122 | 115 | |
| 2907 | 105 | 104 | 104 | 103 | 111 | 93 | 99 | 119 | 97 | 105 | 100 | 109 | 103 | 116 | 1 | 1 | 1 | 1 | 2 | 0 | 3 | 3 | 123 | 117 | |
| 2895 | 105 | 110 | 104 | 108 | 108 | 95 | 99 | 121 | 98 | 103 | 100 | 108 | 105 | 114 | 124 | 118 | 122 | 113 | 117 | 127 | 111 | 120 | 3 | 3 | Group E |
| 2904 | 103 | 107 | 104 | 106 | 108 | 97 | 100 | 123 | 98 | 103 | 100 | 108 | 103 | 113 | 102 | 98 | 95 | 101 | 102 | 106 | 95 | 104 | 2 | 2 | |

C.

PfRh4

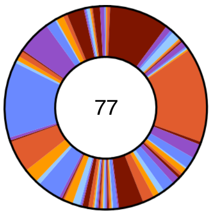


D.

**Competing nanobody**

| Immobilised nanobody | 2858 | 2860 | 2862 | 2863 | 2864 | 2865 | 2867 | 2869 | 2871 | 2872 | 2875 | 2877 | 2859 | 2861 | 2868 | 2870 | 2873 | 2874 | 2866 | 2876 | Group |
|---|---|---|---|---|---|---|---|---|---|---|---|---|---|---|---|---|---|---|---|---|---|
| 2858 | 2 | 3 | 0 | -2 | 2 | 2 | 1 | 1 | 3 | 4 | 4 | 3 | 118 | 111 | 109 | 110 | 118 | 110 | 108 | 115 | Group A |
| 2860 | 4 | 4 | 5 | 3 | 4 | 5 | 4 | 6 | 6 | 8 | 8 | 8 | 116 | 110 | 113 | 110 | 116 | 113 | 109 | 116 | |
| 2862 | 3 | 3 | 3 | 3 | 3 | 4 | 1 | 5 | 4 | 6 | 6 | 6 | 116 | 110 | 114 | 112 | 118 | 115 | 108 | 112 | |
| 2863 | 3 | 3 | 3 | 1 | 3 | 3 | 2 | 3 | 4 | 6 | 5 | 5 | 115 | 109 | 110 | 108 | 115 | 110 | 105 | 112 | |
| 2864 | 3 | 3 | 3 | 0 | 3 | 3 | 2 | 2 | 3 | 5 | 5 | 4 | 115 | 110 | 107 | 106 | 117 | 107 | 105 | 113 | |
| 2865 | 3 | 3 | 0 | -2 | 3 | 2 | 1 | 1 | 4 | 5 | 2 | 3 | 115 | 106 | 102 | 103 | 110 | 99 | 107 | 112 | |
| 2867 | 3 | 4 | 4 | 4 | 4 | 5 | 2 | 7 | 7 | 8 | 8 | 8 | 131 | 122 | 128 | 129 | 130 | 129 | 109 | 130 | |
| 2869 | 2 | 2 | 3 | 3 | 1 | 3 | 2 | 4 | 4 | 5 | 4 | 3 | 118 | 110 | 112 | 112 | 111 | 110 | 110 | 115 | |
| 2871 | 5 | 4 | 3 | 2 | 4 | 5 | 3 | 3 | 5 | 6 | 5 | 6 | 118 | 109 | 110 | 111 | 114 | 110 | 108 | 116 | |
| 2872 | 2 | 2 | -0 | -2 | 1 | 1 | 1 | 1 | 1 | 4 | 2 | 4 | 124 | 117 | 109 | 119 | 121 | 113 | 112 | 118 | |
| 2875 | 3 | 0 | 3 | 0 | 2 | 2 | 1 | 3 | 2 | 5 | 1 | 5 | 116 | 111 | 111 | 111 | 113 | 108 | 107 | 112 | |
| 2877 | 3 | 2 | 3 | 1 | 2 | 3 | 2 | 4 | 3 | 7 | 4 | 4 | 110 | 105 | 108 | 108 | 110 | 106 | 107 | 113 | |
| 2859 | 112 | 77 | 119 | 94 | 111 | 118 | 92 | 82 | 106 | 100 | 95 | 87 | 1 | 4 | 4 | 2 | 6 | 15 | 104 | 118 | Group B |
| 2861 | 97 | 59 | 116 | 80 | 95 | 99 | 85 | 71 | 89 | 102 | 77 | 77 | 5 | 6 | 8 | 3 | 9 | 17 | 102 | 119 | |
| 2868 | 91 | 89 | 124 | 94 | 97 | 91 | 105 | 90 | 105 | 98 | 85 | 95 | 6 | 5 | 9 | 5 | 3 | 16 | 106 | 121 | |
| 2870 | 96 | 78 | 127 | 88 | 82 | 90 | 93 | 87 | 103 | 109 | 73 | 88 | 7 | 3 | 10 | 3 | 3 | 18 | 105 | 116 | |
| 2873 | 113 | 77 | 141 | 104 | 121 | 114 | 108 | 89 | 117 | 118 | 86 | 103 | 3 | 4 | 3 | -1 | 0 | 8 | 109 | 126 | |
| 2874 | 93 | 89 | 171 | 99 | 93 | 91 | 158 | 122 | 126 | 123 | 96 | 124 | 16 | 9 | 14 | 6 | 6 | 22 | 119 | 151 | |
| 2866 | 104 | 102 | 103 | 103 | 109 | 108 | 107 | 103 | 103 | 100 | 105 | 104 | 115 | 111 | 109 | 110 | 114 | 109 | -0 | 3 | Group C |
| 2876 | 112 | 113 | 91 | 115 | 120 | 121 | 113 | 117 | 114 | 115 | 129 | 113 | 125 | 114 | 116 | 113 | 117 | 113 | 3 | 2 | |

E.

PvRBP2b

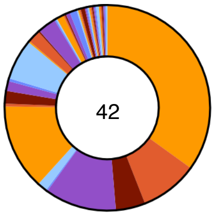


F.

**Competing nanobody**

| Immobilised nanobody | 2910 | 2913 | 2914 | 2919 | 2920 | 2922 | 2924 | 2925 | 2931 | 2934 | 2936 | 2937 | 2923 | 2917 | 2929 | 2930 | 2933 | 2916 | 2932 | 2915 | 2911 | 2927 | 2938 | Group |
|---|---|---|---|---|---|---|---|---|---|---|---|---|---|---|---|---|---|---|---|---|---|---|---|---|
| 2910 | 2 | 2 | 2 | 2 | 2 | 2 | 2 | 2 | 2 | 5 | 1 | 123 | 129 | 126 | 125 | 120 | 136 | 126 | 121 | 123 | 120 | 129 | 123 | Group A |
| 2913 | 3 | 2 | 2 | 2 | 2 | 2 | 3 | 3 | 3 | 7 | 2 | 126 | 130 | 127 | 126 | 122 | 136 | 125 | 123 | 124 | 124 | 130 | 126 | |
| 2914 | 1 | 1 | 1 | 2 | 2 | 1 | 2 | 2 | 2 | 4 | 1 | 123 | 129 | 126 | 124 | 120 | 135 | 124 | 120 | 121 | 121 | 128 | 125 | |
| 2919 | 3 | 2 | 2 | 2 | 3 | 2 | 3 | 3 | 3 | 6 | 2 | 124 | 129 | 128 | 126 | 121 | 136 | 128 | 124 | 125 | 124 | 130 | 126 | |
| 2920 | 1 | 1 | 1 | 1 | 0 | 1 | 1 | 0 | 0 | 3 | 1 | 123 | 128 | 126 | 124 | 120 | 134 | 125 | 120 | 121 | 121 | 129 | 121 | |
| 2922 | 4 | 2 | 3 | 3 | 4 | 2 | 4 | 4 | 3 | 7 | 3 | 126 | 131 | 128 | 127 | 123 | 138 | 129 | 124 | 123 | 125 | 129 | 129 | |
| 2924 | 2 | 2 | 2 | 3 | 2 | 2 | 3 | 3 | 2 | 6 | 2 | 124 | 129 | 127 | 126 | 122 | 136 | 127 | 123 | 122 | 122 | 131 | 97 | |
| 2925 | 2 | 2 | 2 | 2 | 2 | 2 | 3 | 2 | 2 | 5 | 2 | 125 | 130 | 127 | 127 | 123 | 136 | 127 | 121 | 123 | 123 | 129 | 125 | |
| 2931 | 3 | 3 | 3 | 3 | 2 | 2 | 3 | 3 | 3 | 5 | 2 | 127 | 131 | 128 | 127 | 124 | 136 | 128 | 121 | 123 | 123 | 129 | 125 | |
| 2934 | 3 | 3 | 3 | 3 | 4 | 3 | 4 | 3 | 4 | 5 | 3 | 126 | 130 | 127 | 130 | 123 | 137 | 132 | 122 | 124 | 124 | 131 | 128 | |
| 2936 | 1 | 1 | 1 | 1 | 1 | 1 | 1 | 0 | 1 | 4 | 1 | 126 | 130 | 129 | 127 | 122 | 135 | 127 | 122 | 122 | 123 | 131 | 125 | |
| 2937 | 129 | 130 | 130 | 130 | 128 | 130 | 129 | 128 | 133 | 127 | 131 | 17 | 4 | 5 | 4 | 4 | 4 | 3 | 5 | 72 | 87 | 59 | 126 | Group B |
| 2923 | 132 | 129 | 131 | 131 | 133 | 131 | 132 | 132 | 135 | 131 | 134 | 59 | 2 | 8 | 3 | 4 | 2 | 2 | 3 | 123 | 125 | 128 | 125 | |
| 2917 | 129 | 130 | 130 | 131 | 131 | 131 | 130 | 129 | 133 | 130 | 133 | 52 | 2 | 7 | 1 | 2 | 2 | 1 | 1 | 110 | 115 | 109 | 127 | |
| 2929 | 134 | 134 | 135 | 133 | 137 | 135 | 135 | 135 | 136 | 133 | 136 | 72 | 3 | 12 | 2 | 5 | 3 | 2 | 5 | 130 | 130 | 134 | 131 | |
| 2930 | 134 | 133 | 134 | 132 | 135 | 134 | 135 | 134 | 136 | 132 | 136 | 67 | 3 | 11 | 3 | 5 | 3 | 4 | 5 | 128 | 128 | 130 | 131 | |
| 2933 | 129 | 128 | 129 | 129 | 132 | 131 | 130 | 129 | 131 | 128 | 134 | 49 | 2 | 7 | 2 | 3 | 2 | 2 | 1 | 123 | 123 | 127 | 126 | |
| 2916 | 131 | 130 | 131 | 131 | 133 | 133 | 132 | 130 | 135 | 133 | 134 | 77 | 2 | 12 | 2 | 5 | 3 | 3 | 4 | 104 | 125 | 129 | 127 | Group C |
| 2932 | 135 | 135 | 137 | 133 | 137 | 135 | 137 | 136 | 136 | 134 | 136 | 76 | 2 | 13 | 2 | 4 | 2 | 3 | 2 | 70 | 11 | 25 | 133 | Group D |
| 2915 | 131 | 129 | 128 | 129 | 133 | 118 | 132 | 131 | 134 | 131 | 136 | 117 | 118 | 121 | 124 | 120 | 135 | 21 | 5 | 7 | 3 | 3 | 127 | Group E |
| 2911 | 131 | 130 | 129 | 130 | 133 | 131 | 130 | 130 | 133 | 129 | 135 | 126 | 135 | 131 | 129 | 126 | 138 | 119 | 33 | 7 | 2 | 2 | 127 | Group F |
| 2927 | 133 | 133 | 133 | 133 | 136 | 132 | 134 | 132 | 134 | 131 | 138 | 118 | 132 | 128 | 130 | 125 | 139 | 121 | 39 | 10 | 4 | 3 | 5 | Group G |
| 2938 | 130 | 131 | 133 | 131 | 134 | 133 | 100 | 131 | 133 | 132 | 132 | 128 | 131 | 130 | 130 | 126 | 137 | 130 | 125 | 127 | 128 | 132 | 2 | Group H |

Figure S2

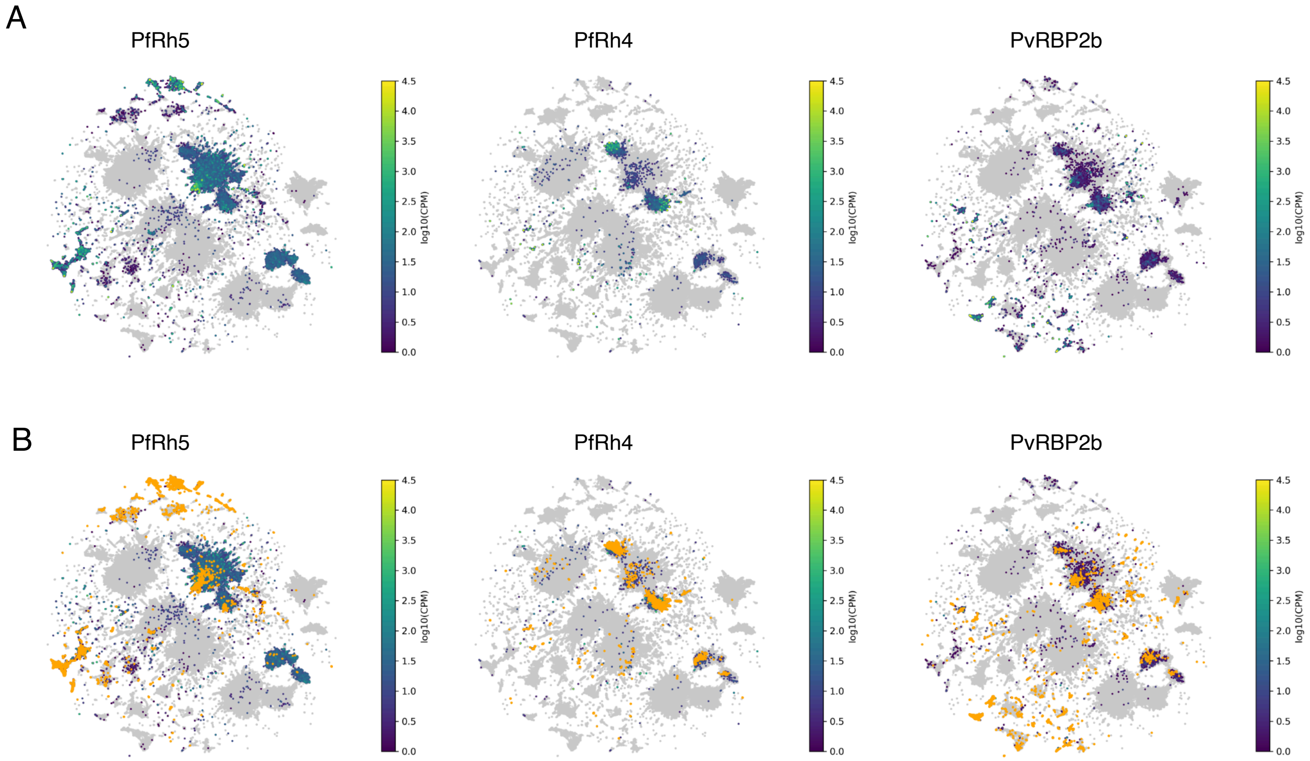
A
PfRh5
PfRh4
PvRBP2b
4.5
4.0
3.5
3.0
2.5
2.0
1.5
1.0
0.5
0.0
log10(CPM)
B
PfRh5
PfRh4
PvRBP2b

Figure S3

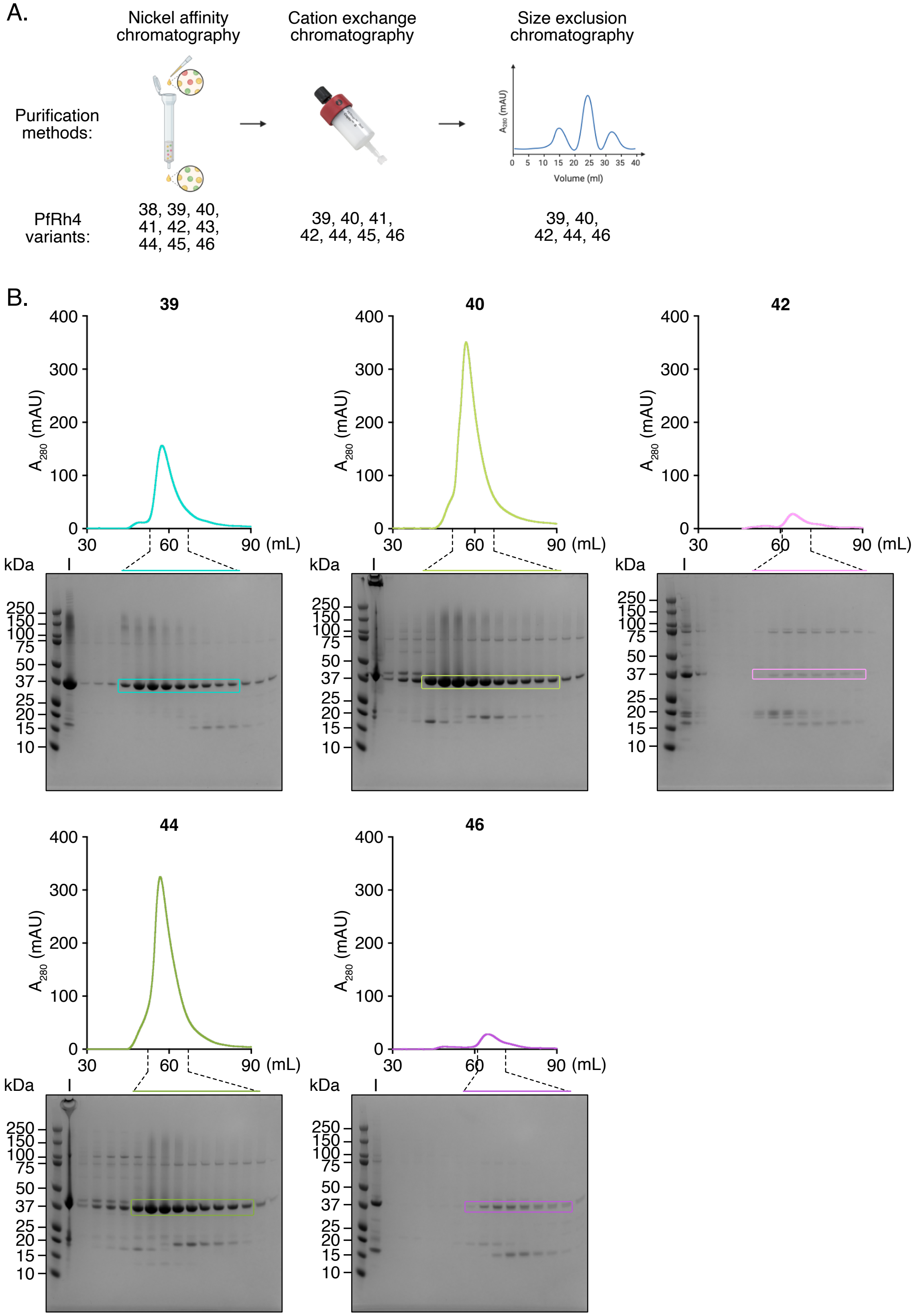
A.
Nickel affinity chromatography
Cation exchange chromatography
Size exclusion chromatography
Purification methods:
A280 (mAU)
Volume (ml)
0 5 10 15 20 25 30 35 40
PfRh4 variants:
38, 39, 40, 41, 42, 43, 44, 45, 46
39, 40, 41, 42, 44, 45, 46
39, 40, 42, 44, 46
B.
39
40
42
44
46
A280 (mAU)
400
300
200
100
0
30
60
90 (mL)
kDa
I
250
150
100
75
50
37
25
20
15
10

Table S1

| | 2739 | 2740 | 2741 | 2742 | 2743 | 2744 | 2745 | 2746 |
|---|---|---|---|---|---|---|---|---|
| 1 | N138D | R139K | M141I | M141I | N138D | M141I | M141I | M141I |
| 2 | R139L | M141I | D177E | C182M | M141I | I197L | C182M | C182M |
| 3 | M141I | D177E | C182M | I197L | L143I | F199L | I197L | I197L |
| 4 | Y142K | S184E | S184E | F199K | T181L | K230I | F199L | F199K |
| 5 | L143I | S195A | I197L | W213Y | C182M | Y244L | K230I | K230I |
| 6 | R167K | I197L | F199L | K230I | I197L | I267L | Y244L | Y244L |
| 7 | D177E | F199L | W213Y | E237I | F199L | S270T | L274F | C276F |
| 8 | T181L | W213Y | K230I | Y244L | K230I | C276F | C276F | V345A |
| 9 | C182M | K230I | M233L | H255R | I234L | I283L | Y279L | Y357F |
| 10 | S184Q | M233K | E237I | C276F | Y244L | V345N | V345C | E393T |
| 11 | N191Q | E237I | Y244L | V345A | I267L | Y357F | Y357F | |
| 12 | S195A | Y244L | H255R | Y357F | S270T | V364I | V364I | |
| 13 | K196N | H255R | L274F | E393T | C276F | E393T | E393T | |
| 14 | I197L | I267L | C276F | K422I | Y279W | Y394I | Y394I | |
| 15 | F199L | S270T | Y279L | | I283L | T403L | T403L | |
| 16 | K200I | C276F | T293S | | H339L | | | |
| 17 | W213Y | I283L | V345C | | V345A | | | |
| 18 | E226L | T293D | Y357F | | S346A | | | |
| 19 | K230I | H339K | V364I | | I353L | | | |
| 20 | M233K | V345N | F381L | | E354L | | | |
| 21 | I234L | Y357F | E393T | | Y357F | | | |
| 22 | E237I | V364I | Y394I | | V364I | | | |
| 23 | Y244L | F381L | T403L | | K366L | | | |
| 24 | K247E | E393T | K422L | | Y378F | | | |
| 25 | H255R | Y394I | | | E393T | | | |
| 26 | I267L | T403L | | | Y394I | | | |
| 27 | S270T | K422I | | | I398L | | | |
| 28 | C276F | | | | T403L | | | |
| 29 | Y279W | | | | | | | |
| 30 | I283L | | | | | | | |
| 31 | T293D | | | | | | | |
| 32 | H339L | | | | | | | |
| 33 | V345A | | | | | | | |
| 34 | S346A | | | | | | | |
| 35 | I353L | | | | | | | |
| 36 | E354L | | | | | | | |
| 37 | Y357F | | | | | | | |
| 38 | V364I | | | | | | | |
| 39 | K366L | | | | | | | |
| 40 | Y378F | | | | | | | |
| 41 | F381L | | | | | | | |
| 42 | E393T | | | | | | | |
| 43 | Y394I | | | | | | | |
| 44 | I398L | | | | | | | |
| 45 | T403L | | | | | | | |
| 46 | S413E | | | | | | | |
| 47 | S417T | | | | | | | |
| 48 | K422I | | | | | | | |
| 49 | Q436K | | | | | | | |
